# Single nucleotide polymorphisms and Zn transport by nuclear ZIP11 shape cancer phenotypes in HeLa cells

**DOI:** 10.1101/2023.08.12.553076

**Authors:** Elizabeth Y. Kim, Odette Verdejo-Torres, Karla Diaz-Rodriguez, Farah Hasanain, Leslie Caromile, Teresita Padilla-Benavides

## Abstract

Zinc (Zn) is an essential micronutrient that regulates critical biological processes such as enzymatic function, gene expression, and cell signaling and provides structural stability to proteins. Under physiological conditions, Zn is a divalent cation (Zn^2+^) in an inactive redox state. Maintaining Zn homeostasis is essential for normal cell development and function, and any dysregulation in supply and transport can lead to pathophysiological conditions. Zn transporters, such as ZIP11, are critical regulators in Zn homeostasis. In mammals, ZIP11 belongs to the GufA subfamily of ZIP transporters and is primarily found in the nucleus and the Golgi apparatus. Our laboratory recently reported an essential role of ZIP11 in maintaining nuclear Zn levels in the cervical cancer cell line HeLa that supports various hallmark phenotypes of cancer. Genomic analysis of publicly available cervical and ovarian cancer patient datasets identified several single-nucleotide polymorphisms (SNPs) in the ZIP11 coding region that correlated with disease severity. We hypothesized that these SNPs might have potentially deleterious consequences as they are in coding regions that could affect ZIP11 function by increasing substrate accessibility, potentially enhancing the carcinogenic phenotype of HeLa cells. In addition, we identified a classic Zn-metal binding site (MBS) composed of three relevant residues which may be required for transmembrane Zn-transport, maintenance of metal homeostasis, and the carcinogenic properties of HeLa cells. To address these questions, we utilized our well-established model of stably knock down (KD) *ZIP11* in HeLa cells and overexpressed ZIP11 encoding single mutations corresponding to the SNPs and the MBS. Overexpression of ZIP11 encoding SNPs restored the Zn levels and the proliferation, migration, and invasive defects of *ZIP11* KD cells. On the other hand, cells expressing ZIP11 with single MBS mutations exhibited a phenotype similar to KD cells, suggesting that Zn transport by this transporter is necessary for establishing and maintaining carcinogenic properties. The work highlights the functional relevance of nuclear Zn transport by ZIP11 to maintain homeostasis and support carcinogenic properties in ovarian cancer cells.

## INTRODUCTION

Zinc (Zn) is one of the most abundant elements in the environment and plays a crucial role as an essential micronutrient. It involves various biological processes, including enzymatic activity, oxidative stress maintenance, transcriptional regulation, signaling, cellular aging, apoptosis, and adequate immune responses^1, 2^. Under physiological conditions, Zn is a divalent cation (Zn^2+^) in an inactive redox state, making it vital for numerous cellular functions^1^. As such, Zn is a cofactor for over 300 enzymes and plays a crucial role in stabilizing the structure of metabolic and signaling enzymes and transcription factors. Maintaining Zn homeostasis is essential for controlling normal cell development and function, and any dysregulation in Zn supply and transport can lead to pathophysiological conditions. For instance, Zn deficiency results in skin-related diseases, hypogonadism, anemia, growth delays, alopecia, chronic inflammation, and deficiencies in immune, hepatic, and mental functions^3–9^. Conversely, excess zinc can be toxic, disrupting the acquisition of other micronutrients like copper^10–12^.

Zinc transporters, specifically the ZnT family (solute-linked carrier 30 or SLC39) and the ZIP family (Zrt, Irt-related proteins, solute-linked carrier 39, or SLC39), are essential for maintenance of Zn homeostasis^1^. ZnT transporters controls Zn efflux, while the ZIP family is responsible for Zn influx across membranes into the cytosol^1^. In mammalian cells, there are nine ZnT exporters (ZnT1-8, 10) and fourteen ZIP importers (ZIP1-14)^1^. The ZIP family of transporters is classified into the I, II, LIV, and GufA subfamilies based on sequence similarities ^13–17^. The only available structure for a ZIP transporter is from *Bordetella bronchiseptica* (BbZIP4). Structurally, the amino- and carboxy-termini domains face the extracellular domain and four Cd^2+^ binding sites were identified in the presence of this ion, confirming previous predictions about ZIP transporter architecture^18^. BbZIP4 presents eight transmembrane (TM) helices, forming a tight bundle, with TM2, TM4, TM5, and TM7 forming an inner bundle, while the others surround them^18^. TM2 contains a 36-amino acid-long domain with a kink linked to a conserved proline (P110), while TM4 and TM5 are bent due to proline residues in the metal-binding sites (MBS)^18^. BbZIP4 structure is distinct to other transporters, as it is symmetric, with TM1-TM3 and TM6-TM8 related by a pseudo-two-fold axis almost parallel to the membrane plane^18^. TM4 and TM5 also seem to be symmetrically related, forming an unusual 3+2+3 TM structure.

ZIP transporters are primarily located in the cell membrane and subcellular compartments and present differential expression according to the cellular developmental stage and needs for the metal^1, 13, 16, 19–23^. However, some transporters, such as ZIP11, which is encoded by the *SLC39A11* gene^17^, is the only ZIP transporter located in the nucleus of epithelial cells of the stomach and colon of mice and in the Golgi apparatus of mammary epithelial cells and the cervical cancer cell line HeLa^24, 25^. The *ZIP11* gene contains metal responsive elements (MREs) targeted by Metal Regulatory Transcription Factor 1 (MTF1) as a response to metal levels^17, 24, 25^. ZIP11 is highly expressed in the testes, stomach, ileum, and cecum, with lower expression in the liver, duodenum, jejunum, and colon. Within the gastrointestinal tract, Zn deficiency modestly downregulates *ZIP11* and induces the expression of *ZIP4* in the stomach, suggesting that ZIP4 might take over Zn absorption from the colon in response to deficiency^25^. Studies on human embryonic kidney (HEK) cells showed that overexpression of *ZIP11*-Flag increased levels of intracellular Zn and decreased cell viability upon incubation in high concentrations of Zn^17^. However, additional *Zip11* siRNA-mediated KD experiments in the murine monocyte macrophage cell line Raw264.7 resulted in reduced intracellular Zn concentrations^17^.

Dysregulation of Zn homeostasis has been implicated in various diseases, including cancer. In this regard, ZIP11 gene association analyses linked somatic changes in the gene variants with patient survival data. For instance, variants like rs8081059 are associated with increased risk of renal cell carcinoma, while rs11871756, rs11077654, rs9913017, and rs4969054 are linked to bladder cancer risk^26^. Additionally, ZIP11 expression is increased in pancreatic adenocarcinoma, colorectal cancer, and breast cancer, while showing a negative correlation with glioma grades^27–29^. However, despite these associations, the detailed biological function and mechanism of ZIP11 in cancer development remain largely unexplored. Experiments from our laboratory functionally characterized ZIP11 in HeLa cells and demonstrated that the transporter regulates nuclear Zn homeostasis, and contributes to cell proliferation, among other carcinogenic phenotypes^24^. Functional characterization and RNA-seq analyses of HeLa cells KD of *ZIP11* evidenced the induction of a senescent state due to an increase in levels of Zn in the nucleus, which resulted in increased sensitivity to Zn stress and impairment of several carcinogenic properties^24^.

Our previous report also identified four single nucleotide polymorphisms (SNPs) that correlated with severity of cervical cancer in patients^24^ (**Fig.1**). In addition, we located three classic Zn^2+^-metal binding sites (MBS) within the transmembrane region of ZIP11, which are considered essential for metal transport^24^ (**Fig. 1**). In this work, we followed up our hypothesis that ZIP11 expressing the SNPs may contribute to the carcinogenic properties of HeLa cells, while mutations on the MBS residues might alter Zn transport, and in consequence, a decrease in their aggressive carcinogenic properties. To test this, we used our well-established short hairpin RNA (shRNA) model against *SLC39A11* to reduce *ZIP11* expression in HeLa cervical cancer cells by targeting the UTR region of ZIP11^24^. We then used a lentiviral vector to overexpress ZIP11 with single mutations at the SNPs and MBS, and functionally characterize the changes in the carcinogenic properties of HeLa cells expressing the mutated transporter. Confocal microscopy confirmed the expression and localization of all the mutants of ZIP11 at the nuclear periphery and Golgi vesicles. Reintroduction of the exogenous ZIP11 expressing the SNPs into the KD cells restored nuclear Zn to normal levels, rescued the proliferation defect and sensitivity of cells to Zn, as well as the functional carcinogenic properties. On the contrary, overexpression of ZIP11 mutated at the MBS failed to rescue the phenotype observed in KD cells. The data supports a model where the SNPs detected in ZIP11 of ovarian cancer patients have a deleterious effect, while alterations in Zn transport impair the establishment of the carcinogenic phenotype in HeLa cells, in a similar manner than the KD of the transporter. Studies to understand the role of Zn transporters, such as ZIP11, and their relationship with metal homeostasis and cancer will provide mechanistic insights of disease progression and potential therapeutic targets.

**Figure 1.**
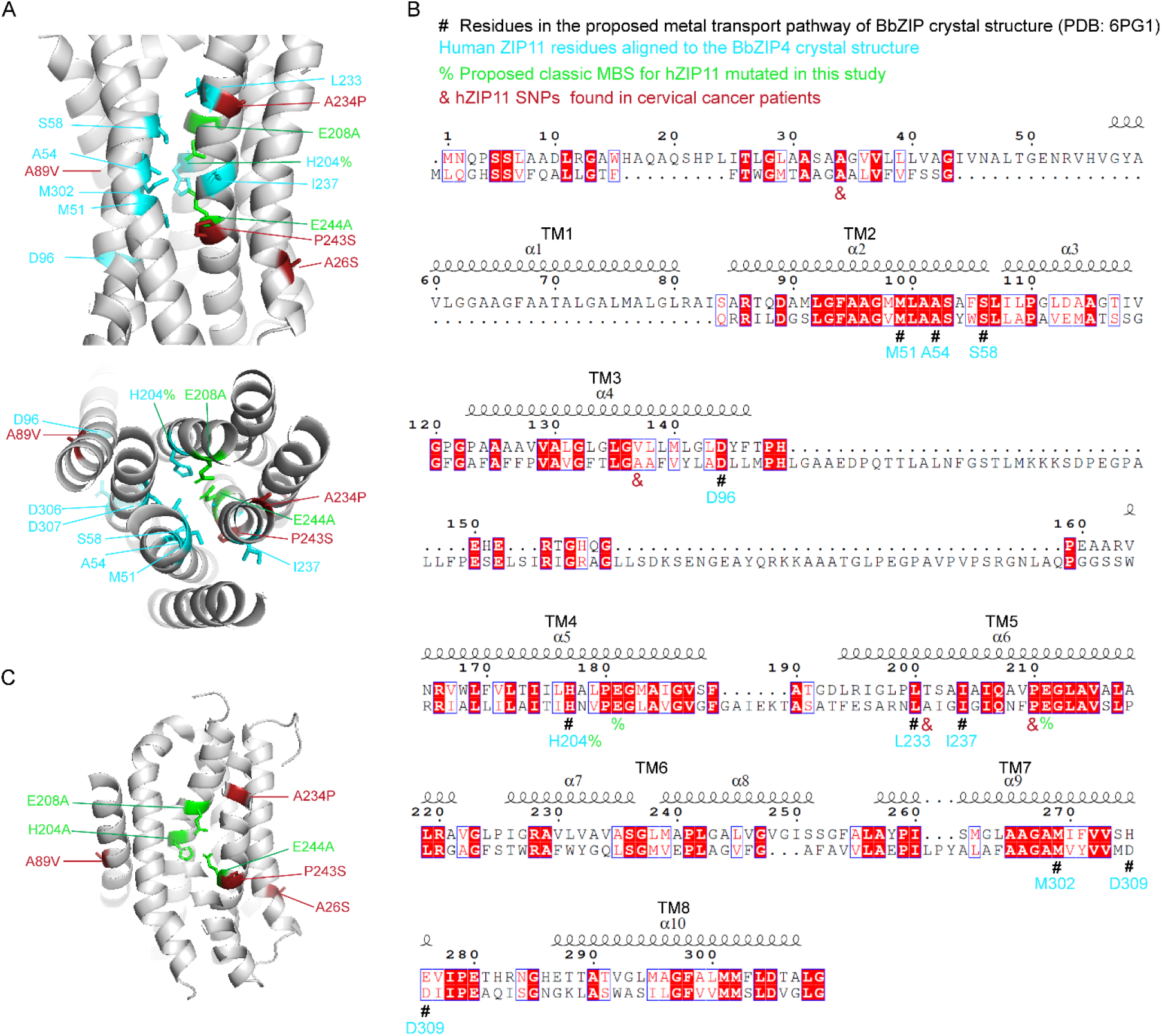
Relevant residues of human ZIP11 compared to BbZIP4. **A.** Alpha Fold predicted model of BbZIP4 and human ZIP11 with SNPs (red), MBS mutations (green) and additional residues located in the proposed pathway of metal transport in BbZIP4 (blue) labeled. **B.** Alignment of SNPs and MBS of BbZIP4 (top) compared to the human ZIP11 (bottom). Residues of human ZIP11 aligned to residues of the BbZIP4 crystal structure are labeled in blue. Other residues involved in the proposed metal transport pathway of BbZIP4 are labeled with a “#”. The TM helices are labeled. **C.** Zoom into the transmembrane region of ZIP11 showing the localization of MBS mutations (green), SNPs (red).

## RESULTS

### Relevant SNPs of ZIP11 in ovarian and cervical cancer and importance of metal transport

Our published analysis of the TCGA database found that uterine corpus endometrial carcinoma patients have the highest incidence of ZIP11 mutations^24^. ZIP11 loss was common in esophageal carcinoma, while increased expression occurred in ovarian cystic adenocarcinoma, breast invasive carcinoma, lung squamous cell carcinoma, bladder urothelial carcinoma, and cervical cancer patients^24^. Elevated ZIP11 expression correlated with poor prognosis in cervical cancer, verified in HeLa cells overexpressing wild type ZIP11^24^. The SNP mutations A26S, A89V, A234P, and P243S were identified, with A234P and P243S predicted to have the strongest impact in disease progression. Bioinformatics analyses hinted that A234P and P243S could alter ZIP11 function in metal transport^24^. On the other hand, A26S and A89V were predicted to be less likely to affect substrate accessibility and Zn transport^24^. These observations led us to the hypothesis that SNPs present in ZIP11 in cancer patients might enhance ZIP11 function contributing to the aggressive cancer phenotype and poor prognosis observed, potentially related to gain of function of ZIP11.

We also identified three MBS residues that have been implicated in metal transport, H204, E208 and E244. Homology modeling and sequence analyses indicated a high conservation and a similar localization of these residues with those proposed in the transport pathway of the ZIP4 transporter from *B. bronchiseptica* BbZIP4^18^, suggesting that the residues in ZIP11 may have a similar function than their counterpart on BbZIP4 (**Fig. 1**). For instance, the H204 in human ZIP11 corresponds to H177 in BbZIP (labeled “#”), and E208 and E244 aligned with the metal-binding residues E181 and E211 in BbZIP4, respectively. The analyses support the established model of the contributions of these canonical residues in releasing metal from the transmembrane transport pathway to the cytosol (**Fig. 1**)^18^.

To understand the biological relevance of the SNPs identified in cervical cancer patients (A26S, A89V, A234P, and P243S), as well as the Zn transporting properties of ZIP11 (H204A, E208A and E244A), we utilized the ZIP11 KD HeLa cells from our previously reported work as a model^24^. Consistent with our previous work, western blot analyses of ZIP11 expression showed that KD cells have a significant reduction in the expression of the transporter compared to wild type control cells, which was restored upon expression of wild type ZIP11 (**Fig. 2A**). ZIP11 encoding the SNPs and the MBS mutants also showed an increased expression compared to KD and control cells (**Fig. 2B**). However, H204A and E208E had a lower expression compared to the rest of the mutants (**Fig. 2B**).

**Figure 2.**
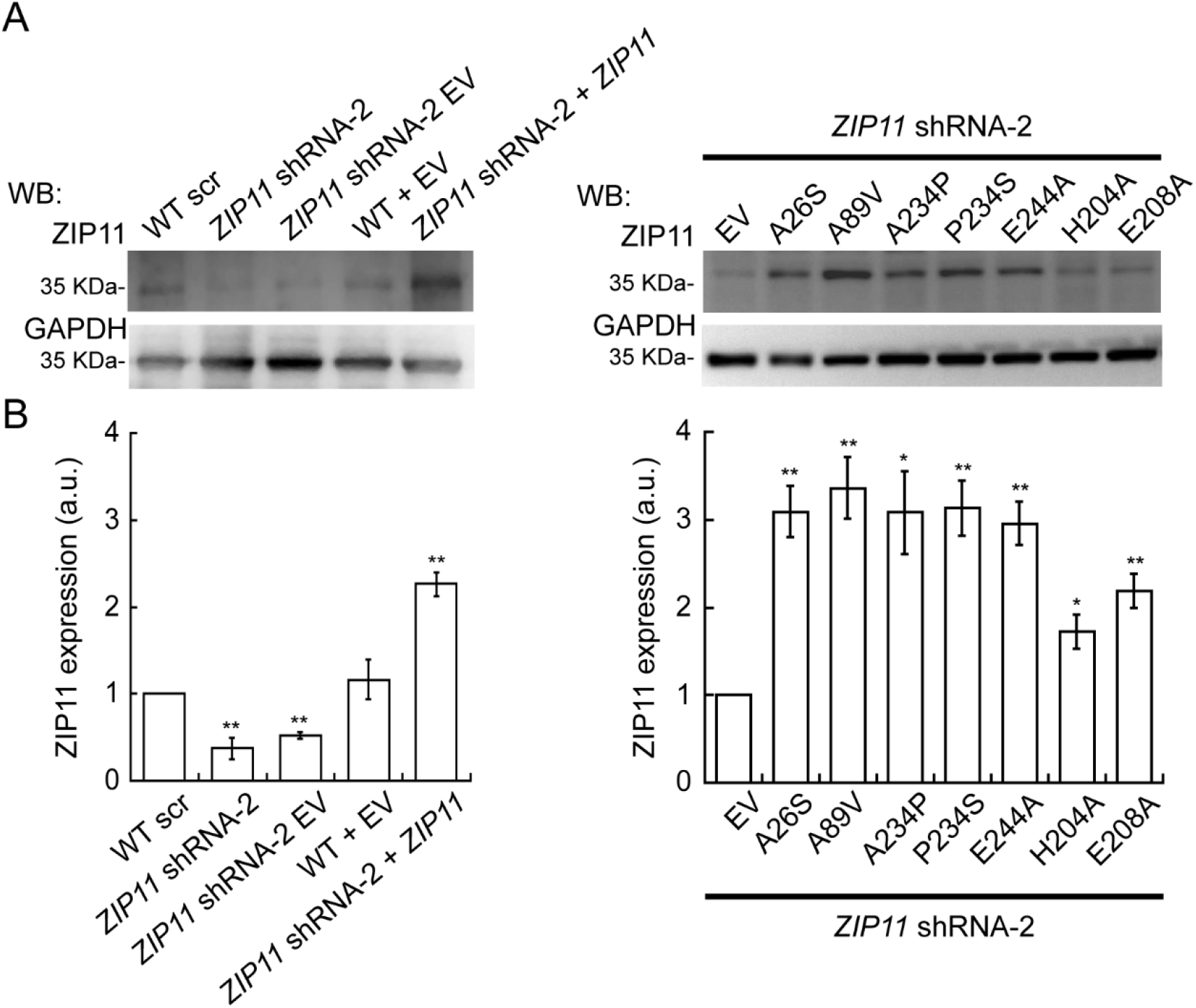
Expression of SNPs and MBS mutants in HeLa cells KD for *ZIP11.* **A.** Representative western blot of ZIP11 in proliferating wild type (WT) and *ZIP11* KD HeLa cells transduced with control vectors and a lentiviral vector encoding for *ZIP11* wild type, the four SNPs (A26S, A89V, A234P, and P243S) and the three MBS (H204A, E208A, E244A) mutations. **B.** Quantification of *ZIP11* expression levels in HeLa cells shown in A. N=4; *P < 0.05, **P < 0.01.

To verify this phenotype and determine the localization of these proteins, we performed confocal microscopy analyses. We found that the mutant ZIP11 proteins localized predominately to the vesicles as compared to the controls; however, in all cases these vesicles were also located at the periphery of the nuclei (**Fig. 3**). Consistent with the western blot analyses, the expression of the four SNPs (A26S, A89S, A234P and P243S) and the E244A mutant was elevated in KD cells (**Fig. 3A, B**), while the H204A and E208A were less abundant (**Fig. 3B**). The green fluorescent protein (GFP) was used as a control of transduction efficiency, as the pLV plasmid encodes the *GFP* and can be used as a proxy measure of successful integration of the GFP and ZIP11 cassettes in the genome of our KD HeLa cells (**Fig. 3**, cyan panel).

**Figure 3.**
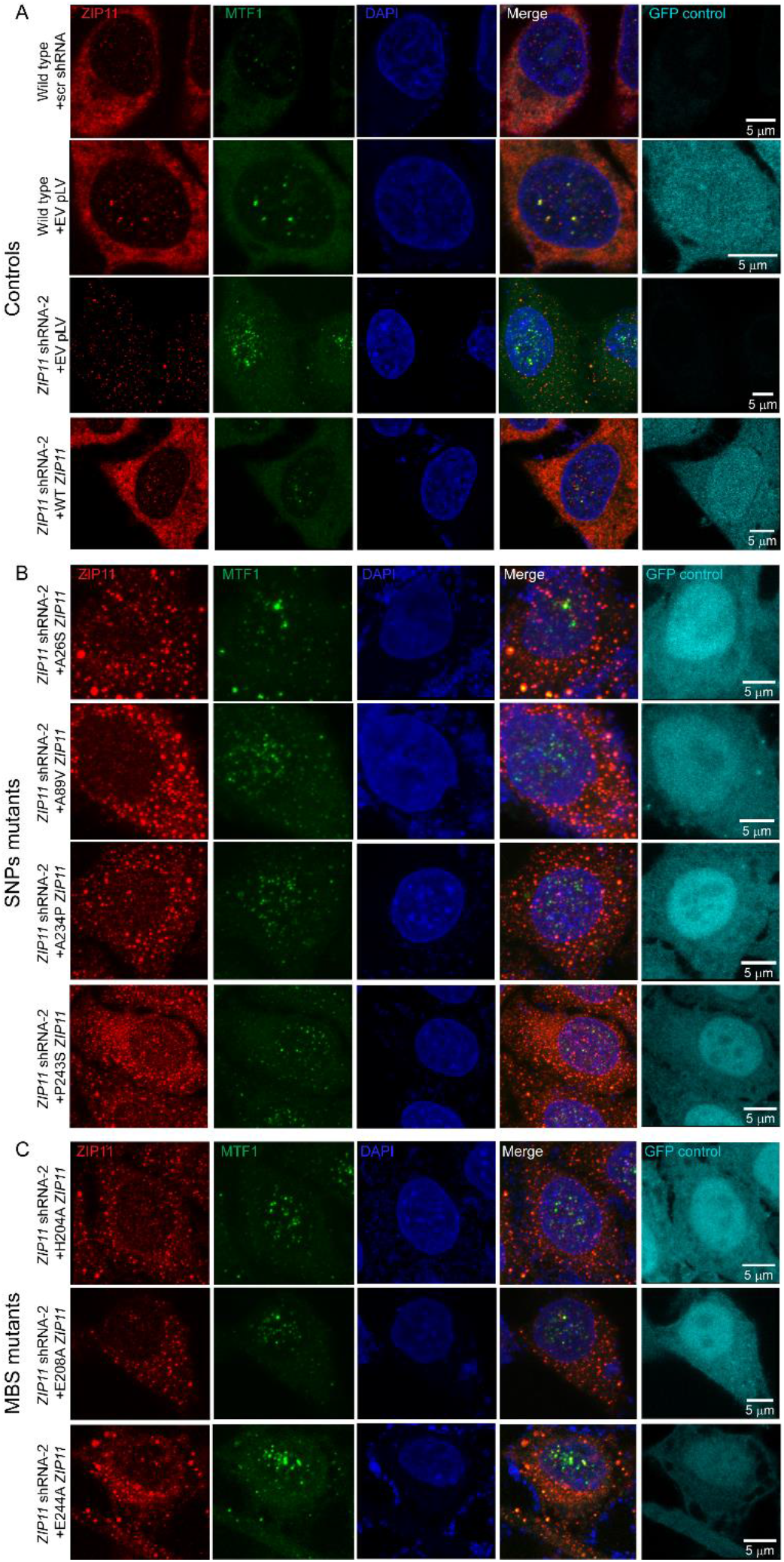
ZIP11 localization in HeLa cells. Representative confocal microscopy images of ZIP11 (red), MTF1 (green) and nuclei stained with DAPI (blue). GFP was used as a control for transduction. **A.** Confocal images show a decrease in staining for ZIP11 in HeLa cells transduced with *ZIP11* shRNA-2 and the pLV1 empty vector compared to wild type cells transduced with scr shRNA, or the pLV empty vector. Expression is recovered in KD cells reconstituted with exogenous ZIP11^24^. **B.** Confocal images of *ZIP11* KD cells expressing the SNPs A26S, A89V, A234P, and P243S show a restoration in staining in the nuclear and cytosolic regions. **C.** Confocal images of *ZIP11* shRNA cells expressing the MBS mutations H204A, E208A, and E244A show reduced staining for ZIP11. Bar = 5 μm.

### Nuclear expression MTF1 is slightly elevated in cells expressing ZIP11 encoding the MBS mutations

We have previously shown that the expression of MTF1 protein was elevated in cells KD for *ZIP11* and restored to basal levels in cells reconstituted with the wild type *ZIP11* gene (**Fig. 3**)^24^. Thus, we sought to determine whether the expression of the SNPs or MBS mutations in the transporter would trigger the expression of this metal-homeostasis related transcription factor. We found that KD *ZIP11* cells expressing the SNP and MBS mutants exhibited decreased MTF1 expression, but not in the cells expressing the MBS mutations alone, suggesting that metal homeostasis remains altered in a similar manner that KD cells (**Fig. 3**).

### SNP mutations, but not MBS mutations, restore nuclear Zn levels in KD *ZIP11* cells

Our previous studies showed that KD *ZIP11* HeLa cells accumulated nuclear Zn, indicating that the transporter is necessary to maintain nuclear levels of the metal^24^. To understand the effect of the SNPs and mutations of a classic transmembrane MBS on the ability of ZIP11 to mobilize Zn from the nucleus, we obtained subcellular fractions from control HeLa cells and those transduced with the ZIP11 mutants and determined the metal concentrations by atomic absorption spectroscopy (AAS, **Fig. 4**). Control *ZIP11* KD cells and those reconstituted with the empty vector had significantly higher levels (two fold) of nuclear Zn compared to wild type controls, which were restored by introducing the exogenous wild type *ZIP11* gene, as previously described (**Fig. 4A**)^24^. We then evaluated the effect of SNPs on the nuclear distribution of Zn. The A26S, A234P and P243S mutations resulted in reduced levels of nuclear Zn similar to that of wild type controls, indicating that these SNPs may contribute to nuclear Zn balance. However, cells expressing the SNP A89V maintained significantly higher levels of nuclear Zn, similar to that of the ZIP11 KD cells (**Fig. 4A**), suggesting that this residue may negatively impact ZIP11’s ability to mobilize Zn out of the nucleus. We then analyzed the contributions to Zn transport of individual MBS that are located at the interphase of the membrane and are essential to enable translocation of the metal. As expected, *ZIP11* KD cells expressing the transporter encoding the H204A, E208A and E244A mutations retained elevated levels of nuclear Zn similar to the KD cells. In terms of cytosolic Zn content, we detected a small non-significant increase in all the strains tested (**Fig. 4B**). Only the introduction of the MBS E208A mutant resulted in a significant cytosolic increase of Zn (**Fig. 4B**). Consistently, analyses of whole cell levels of Zn were elevated in many cases, but only were significant for the A89V, A234P, E208A and E244A mutants (**Fig. 4C**). Based on the location of these residues, the data agrees with the model of dynamic changes of Zn levels in the nuclear fraction according to the potential accessibility and mobility of the substrate through the transmembrane domain.

**Figure 4.**
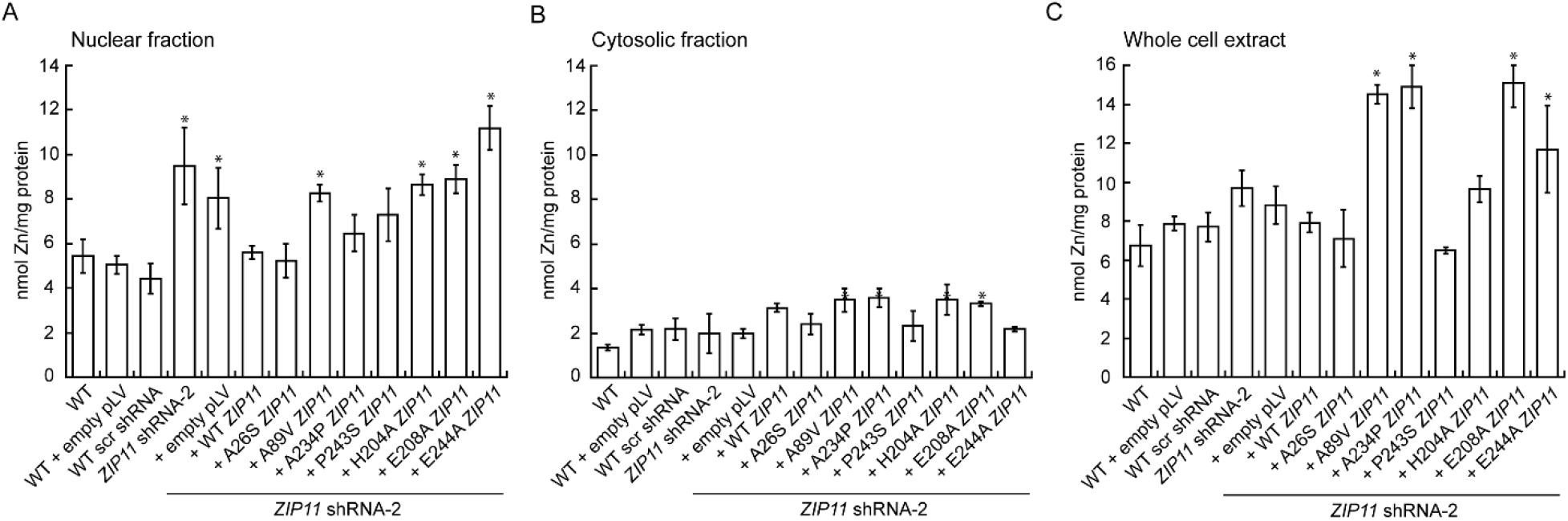
Zn levels in proliferating HeLa cells expressing ZIP11 encoding the SNPs and MBS mutants. WT HeLa cells, WT cells transduced with the *ZIP11* empty vector, scrambled shRNA (scr shRNA), or *ZIP11* shRNA-2, and *ZIP11* shRNA-2 cells expressing the empty vector, exogenous *ZIP11,* SNPs, or MBS mutations were allowed to proliferate for 48 h. Subcellular fractions were obtained and AAS was used to determine the nuclear, cytosolic, and whole cell levels of Zn. **A.** Nuclear fractions. **B.** Cytosolic fractions. **C.** Whole cell extract. N=3; *P < 0.05.

### ZIP11 encoding SNPs detected in cervical cancer patients restore the proliferation defect observed in *ZIP11* KD HeLa cells, while mutations of the MBS are unable to rescue the phenotype

We recently showed that ZIP11 is required for the establishment and maintenance of several hallmark properties of the cervical cancer cell line HeLa^24^. *ZIP11* KD impaired proliferation, migration, and invasion in cultured HeLa cells, and resulted in a senescent state which were rescued by reintroducing the exogenous wild type ZIP11 gene into the KD cells^24^. Therefore, we expanded our studies to understand the functional effect of the four identified SNPs in cervical cancer patients, as well as the effect of essential MBS required for metal mobilization. First, we evaluated the effect of ZIP11 encoding the four SNPs and three MBS mutants on the proliferation of HeLa cells KD for *ZIP11.* Control experiments of KD *ZIP11* cells and those transduced with the empty pLV vector proliferated at a significantly lower rate than wild type cells, recapitulating our previous observations (**Fig. 5A**)^24^. Reconstitution of KD cells with wild type ZIP11 rescued the growth defect, confirming that the transporter is required for the proliferation of HeLa cells, as previously described by our group (**Fig. 5B**)^24^. Analyses of *ZIP11* KD cells transduced with the SNPs A26S, A89V, A234P, and P243S rescued the growth defect, such as these proliferated at a similar rate to that of the control wild type cells and KD cells reconstituted with the wild type transporter (**Fig. 5C**). The data suggest that the SNPs may be functionally relevant to cell growth. On the contrary, cell proliferation assays of cells expressing single mutations of the three MBS in the transporter showed that the proliferation defect observed in the *ZIP11* KD HeLa cells was maintained upon expression of the MBS mutations E208A and H204A. However, the E244A mutant partially restored the growth phenotype (about 40% of the KD cells), but not to the levels of control cells (**Fig. 5D**).

**Figure 5.**
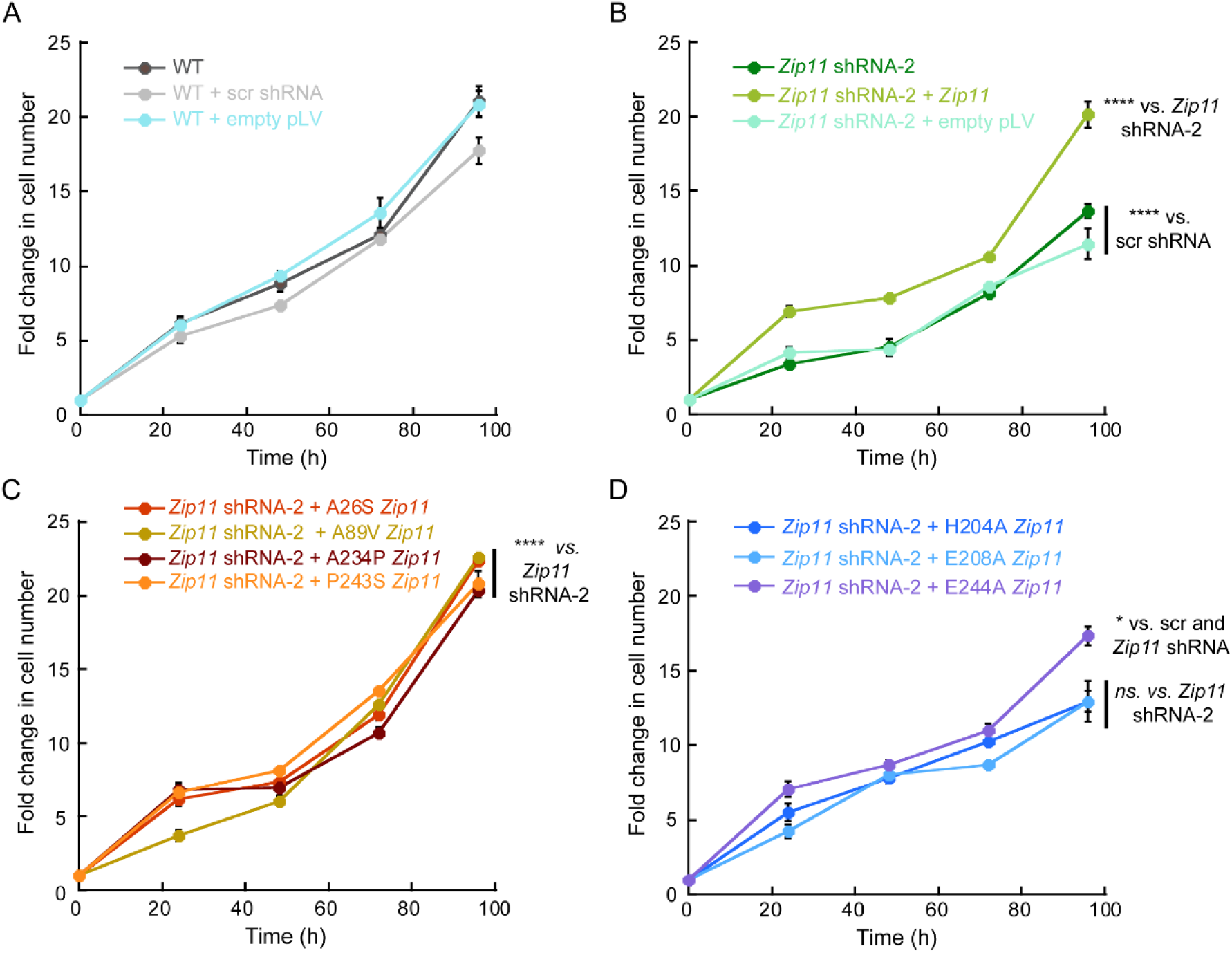
The ZIP11 SNP linked to cervical cancer effectively rescues the proliferation deficiency observed in *ZIP11* KD cells, while the expression of MBS mutations underscores the critical role of metal transport in facilitating cellular growth. Cell counting assay of **A.** proliferating WT, cells transduced with scr shRNA, or the *ZIP11* empty vector (empty pLV); **B.** proliferating cells transduced with *ZIP11* shRNA-2, or *ZIP11* shRNA-2 expressing exogenous *ZIP11* or the empty vector; **C.** proliferating cells transduced with *ZIP11* shRNA-2 expressing the SNPs A26S, A89V, A234P, and P243S. **D.** proliferating cells transduced with *ZIP11* shRNA-2 expressing the MBS mutants H204A, E208A, and E244A grown in normal culture media. N=3; *P < 0.05, ****P < 0.0001.

We previously demonstrated that *ZIP11* KD increased the sensitivity of HeLa cells to elevated levels of Zn in the culture media, a phenotype that was rescued by reintroducing wild type ZIP11 into the KD strain (**Fig. 6A,B**)^24^. Therefore, we investigated the effect of introducing SNP mutations, or whether altering the metal binding capabilities of ZIP11 would have an effect in the Zn resistance phenotype. To this end, the cells were cultured for 72 h with media supplemented with increasing concentrations of Zn. This time point was selected, as our previous work showed that it was the earliest time point where a different behavior was evidenced among the cell lines tested^24^. Cell counting assays showed that the metal sensitivity observed in HeLa cells KD for *ZIP11* was rescued by the transporter encoding the SNPs, however the effect of each mutant was distinctive. For instance, the A26S, A234P, and P243S mutations exerted the largest rescue effect against Zn toxicity. The A26S and A234P reached an increase in cell numbers similar to wild type controls (**Fig. 6C**), while the P243S resulted in a slightly lower survival (50%) at the highest concentration of Zn tested, 200 µM. The A89V mutant restored only 20% of survival rate to Zn stress, compared to wild type controls. On the other hand, ZIP11 encoding the single mutations of the three MBS had a minimal effect in restoring metal resistance (**Fig. 6D**). In this case, expression of ZIP11 encoding the H204A and E208A mutations resulted in an approximately 20% increase in cell survival at high concentration of the metal (200 µM), while the E244A mutant showed an increase of approximately 50% of surviving cells upon metal stress. None of the MBS mutants reached the level of resistance to elevated Zn observed in control cells. These findings suggest that the MBS mutations, especially E244A, might have some impact on the functionality of ZIP11, although to a limited extent.

**Figure 6.**
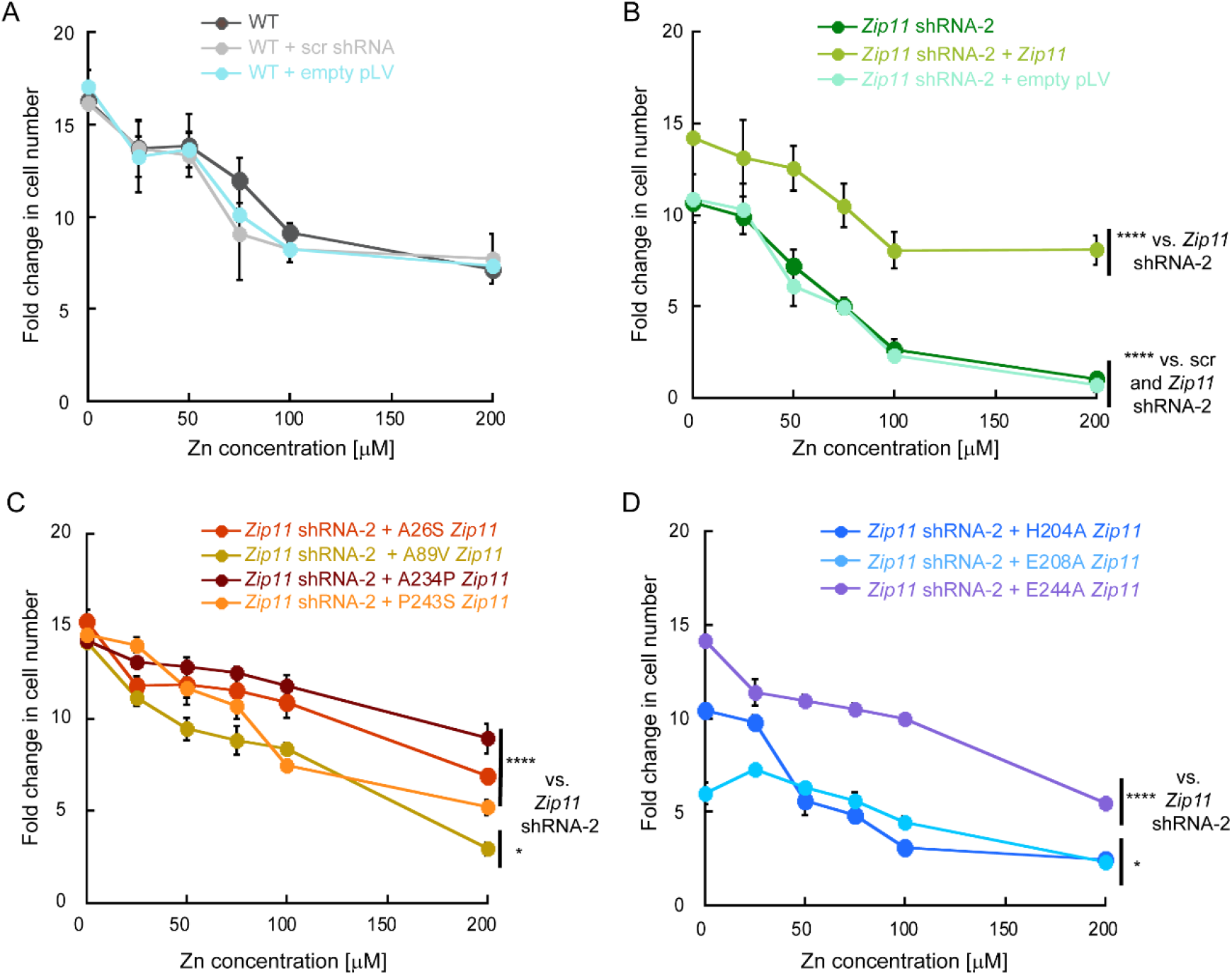
Expression of SNPs in ZIP11 KD HeLa cells restore the sensitivity to Zn while the MBS mutants sustain this heightened sensitivity. Cell counting assay of **A.** proliferating WT, cells transduced with scr shRNA, or the *ZIP11* empty vector (empty pLV); **B.** proliferating cells transduced with *ZIP11* shRNA-2, or *ZIP11* shRNA-2 expressing exogenous *ZIP11* or the empty vector; **C.** proliferating cells transduced with *ZIP11* shRNA-2 expressing the SNPs A26S, A89V, A234P, and P243S. **D.** proliferating cells transduced with *ZIP11* shRNA-2 expressing the MBS mutants H204A, E208A, and E244A cultured for 72 h with increasing concentrations of ZnSO4. N=3; *P < 0.05, ****P < 0.0001.

### ZIP11 encoding SNPs detected in cervical cancer patients, but not the transporter mutated in MBS, restore functional defects associated with fundamental carcinogenic properties *ZIP11* KD HeLa cells

We have reported that *ZIP11* KD HeLa cells have impaired migration and invasive properties, because of nuclear Zn accumulation, and the induction of a senescent state associated with the absence of the transporter^24^. Thus, to assess the effect of the SNPs in ZIP11 on these hallmark carcinogenic properties of HeLa cells, we performed functional assays that assess migration and invasive properties of cancer cells.

First, we performed wound-healing assays, where a scratch was made on a confluent monolayer of cells with a sterile pipette tip and we determined the time taken to close the wound^24, 30–32^. Cell mobility into the area of the wound was monitored at various time points (0, 24, 48, 72, and 96 h) after scratching the monolayer of control and ZIP11-expressing HeLa cells (**Fig. 7**). Time 0 h indicates the moment when the wound was first performed. The area that the cells migrated over time was quantified to determine the rate of wound closure of each cell type. To prevent cell proliferation, before initiating the experiment, the cells were serum-starved overnight and treated with AraC for 2 h to ensure that the observed phenotypes were due to cell mobilization rather than cell division. Consistent with the published data, cells KD for *ZIP11* migrated at a slower rate and covered only 60% of the wound compared to the controls (**Fig. 7A, B**). Reconstitution of wild type ZIP11 in KD cells fully restored the deficient migration (**Fig. 7A, B**)^24^. *ZIP11* KD cells transduced with the SNPs A26S and A234P closed the wound at a similar rate to the control cells (**Fig. 7C, D**), suggesting that these SNPs may restore the migration phenotype associated with ZIP11 function. The P243S mutation had a significantly slower migration rate than control cells and partially restored (80%) the migration capabilities of KD cells, while the A89V SNP presented a similar behavior to the KD cells, indicating that the mobility defect is maintained, as no phenotype recovery was achieved (**Fig. 7C, D**).

**Figure 7.**
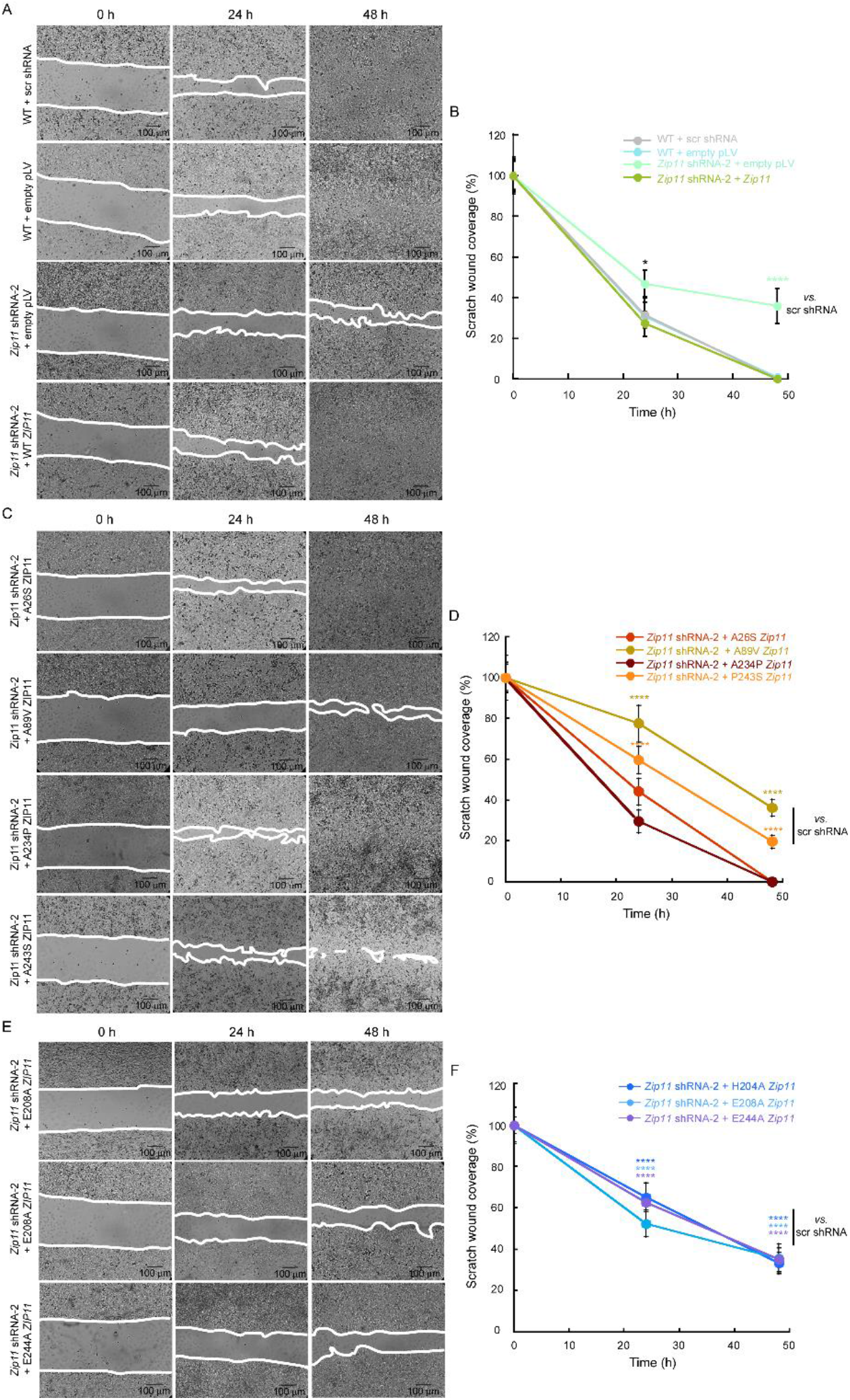
Expression of ZIP11 encoding SNP mutations restores the migration defect in *ZIP11* KD cells, whereas the MBS mutations do not yield the same restorative effect. Representative light microscopy images show the closure of a wound made on a confluent monolayer of cells over 3 days. Time 0 shows the wound immediately after it was inflicted. Wounds were monitored until closure at 48 h. Scale bar = 100 µm. **A.** Wound healing assay and quantification of the area migrated by the cells into the wound over time. **A & B** Control HeLa cells. C & D. *ZIP11* KD cells expressing ZIP11 encoding the SNPs mutations. **E & F**. *ZIP11* KD cells expressing ZIP11 encoding the MBS mutations. *P < 0.05, ****P < 0.0001.

To determine the biological relevance of the residues involved in metal transport, we then tested the effect of MBS mutations on the migration capabilities of HeLa cells. Consistent with the significance of these residues in metal transport, wound healing assays showed that *ZIP11* KD cells expressing the transporter encoding the MBS mutations H204A, E208A and E244A presented a significantly slower migration rate than that of the wild type controls and KD cells reconstituted with normal *ZIP11* (**Fig. 7E, F**). These data strongly suggest that compromised Zn transport also directly or indirectly alters pathways that drive the directional migration of cells.

To assess the contributions of the relevant SNPs and metal transporting properties of ZIP11 on the invasive properties of HeLa cells, we assessed the ability of cells expressing ZIP11 encoding these mutations to invade through the basement membrane substrate Matrigel. Prior to the assay, proliferation was inhibited by serum starvation and AraC treatment. Then, the cells were seeded on invasion chambers with membranes containing 8 µm pores and covered with Matrigel and incubated for 24 h. Consistent with the recently published data, *ZIP11* KD cells did not cross the membrane, while control cells did (**Fig. 8A,B**)^24^. KD cells reconstituted with exogenous ZIP11 completely restored the deficient invasion phenotype (**Fig. 8A,B**)^24^. Similarly, *ZIP11* KD cells expressing the SNPs A26S, A89V, A234P, and P243S also restored the invasion phenotype suggesting that ZIP11 encoding these cancer-related mutations may confer invasive capabilities to the cells (**Fig. 8C,D**)^24^. Conversely, the cells expressing the transporter with MBS mutations were unable to regain the invasive phenotype (**Fig. 8E,F**)^24^, as these cells were unable to invade through the membrane compared to the wild type controls. The data indicates that proper metal binding and transport may be required to enable migration and invasion of HeLa cells. Together, the data confirmed that ZIP11 regulates nuclear Zn homeostasis, and contributes to the establishment of invasive and migration carcinogenic properties of cultured cervical cancer cells.

**Figure 8.**
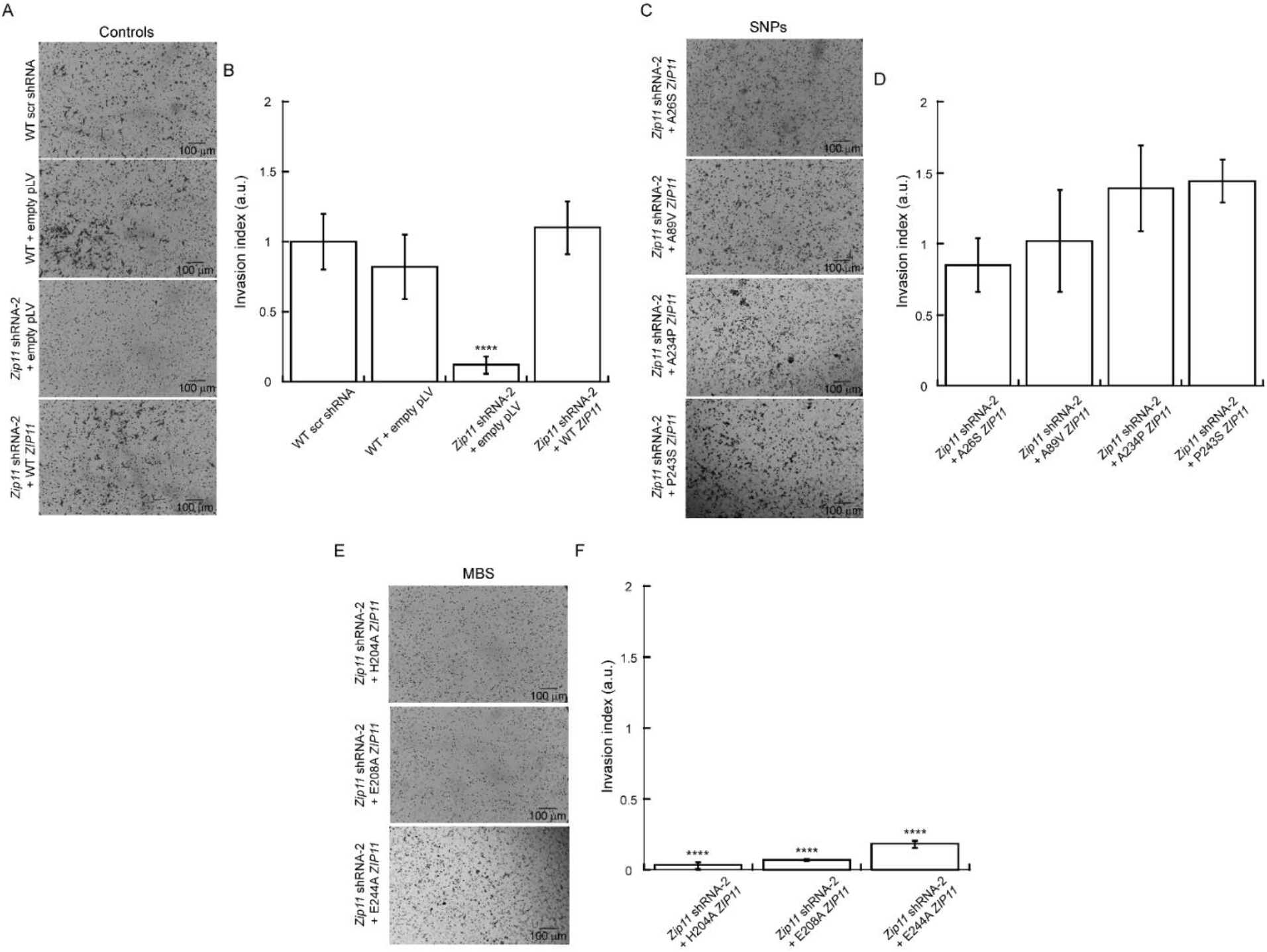
Expression of *ZIP11* encoding the SNPs restored the invasive defect in KD cells, while the mutated MBS resulted essential for this process. Transwell invasion assay and quantification of invading cells. Representative light microscopy images show cells that invaded through Matrigel after 24 h. **A & B** Control HeLa cells. C & D. *ZIP11* KD cells expressing ZIP11 encoding the SNPs mutations. **E & F**. *ZIP11* KD cells expressing ZIP11 encoding the MBS mutations. ****P < 0.0001.

## DISCUSSION

Zinc transporters are key regulators of zinc homeostasis in eukaryotes. Most ZIP transporters are localized to the cell membrane and import extracellular Zn into the cytosol. Some ZIP transporters have specific subcellular localizations which are required for maintenance and appropriate distribution of Zn into organelles, such as ZIP11. The structure of the prokaryotic BbZIP4 provided important insights into the potential mechanism of Zn transport in its eukaryotic homologs, and other ZIP transporters^18^. The structure confirmed a transmembrane domain comprised of eight helices, with specific residues involved in metal binding and transport^18^. The transport pathway involves the recruitment of Zn to the entrance cavity by a conserved serine residue, followed by a conformational change to expose the metal to a chain of metal-binding residues, in a electrogenic, non-saturable, and voltage-dependent manner^18^. The proper function of ZIP transporters is essential for cell viability. As such, alterations of cellular levels of Zn have been implicated in carcinogenesis, due to its role in cell signaling, immune system response, oxidative stress, and apoptosis. In this work, we built upon our previous work on ZIP11, which regulates Zn transport from the nucleus and Golgi into the cytosol of mammalian cells. We demonstrated previously that KD of *ZIP11* results in elevated nuclear Zn levels, which had reduced tolerance to Zn stress, impaired cell proliferation and other carcinogenic properties of HeLa cells.

Cancer cells are characterized by their ability to sustain growth-promoting signaling and activate invasion and metastasis^33^. Understanding the role of Zn transporters, including ZIP11, and their relationship with Zn homeostasis and cancer provides valuable insights into the mechanisms underlying cellular processes and potential therapeutic targets. In this regard, we showed that ZIP11 is required to maintain carcinogenic properties of cervical cancer cells through its regulation of nuclear Zn homeostasis^24^. Here, we explored the functional contributions of specific SNPs residues, identified in cervical cancer patients, and classic amino acids responsible for Zn transport by ZIP11, building upon our observations of the crucial role of the transporter in the carcinogenic properties of HeLa cells^24^. Introducing the identified ZIP11 SNPs (A26S, A89V, A234P, and P243S) into KD cells led to the restoration of deficient proliferation, migration, and invasion phenotypes, implying the promotion of aggressive cancer phenotypes in cervical cancer cells. On the other hand, KD cells expressing ZIP11 MBS mutations (H204A, E208A and E244A) accumulated nuclear Zn, which represents a proxy measure of the impaired metal-transporting function. It is noteworthy that the expression of ZIP11 encoding the MBS mutations was reduced, suggesting that proteins unable to transport Zn might be unstable and degraded at a faster rate than wild type proteins. This data supports the classic model where proper metal binding at these sites in the ZIP11 transport pathway is essential for specific cellular functions.

### The crucial role of Zn in cell proliferation and establishment of carcinogenic properties

Intracellular Zn balance is essential regulating cell proliferation and cell cycle progression. Zn deficiency impairs DNA synthesis and delay cell cycle progression, by affecting the expression of key genes, such as deoxythymidine kinase^34–36^. Increased levels of labile cytosolic Zn has been associated with induction of DNA synthesis and cell proliferation^37^. In this context, our recent research shed light on the significance of ZIP11 in accumulation of nuclear Zn in HeLa cells which delayed proliferation and cell cycle progression^24^. Here, we showed that cells expressing SNPs identified in cervical cancer patients are capable of restoring ZIP11 function and recover the proliferation defect produced by the decreased expression of the transporter in KD cells. In addition, we determined the relevance of three MBS located at the interphase of the membrane. Consistent with previous work, the H204, E208 and E244 residues are essential for metal translocation, which in our model, leads to maintenance of adequate levels of Zn to enable normal cell cycle progression and proliferation of HeLa cells. Our previous RNA-seq analyses, pointed towards potential mechanisms by which the proliferation defect in KD cells could be explained^24^. For instance, cyclin dependent kinase 20 (CDK20), responsible for cyclin-dependent kinase 2 (*CDK2*) activation, was downregulated. As CDK20-mediated phosphorylation of CDK2 is essential for HeLa cell growth, this downregulation likely contributes to the observed growth defect in KD cells^38, 39^. Additionally, *CDKN2C*, a regulator of G1 phase arrest, and *PPP2CA*, encoding the catalytic subunit of PP2A, was upregulated, suggesting a negative control of cell growth in KD cells^39–41^.

The ability to activate invasion and migration are crucial components of metastasis^33^, which results in the spreading of cancer cells from their origin at the primary tumor to a distant location in the body^42^. Our lab previously demonstrated that *ZIP11* KD resulted in migration and invasion defects, indicating that *ZIP11* is required for these metastatic properties^24^. A similar behavior was also observed here in KD cells expressing MBS mutations, supporting the functional role of the transporter in disease progression. The mechanism by which Zn dysregulation due to MBS mutations and ZIP11 KD impairs cell migration and invasion is not yet known. However, evidence from other transporters may shed light into this question. For instance, ZIP13 promote ovarian cancer cell migration and invasion through the activation of the Src/focal adhesion kinase (FAK) signaling pathway^43^. *ZIP13* knock-out altered the subcellular distribution of Zn, and reduced cell migration and invasion due to the decrease in Src and FAK phosphorylation^43^. FAK is a non-receptor tyrosine kinase that is upregulated in ovarian cancers and contributes to cell migration and invasion by facilitating signaling between the cell and extracellular matrix upon phosphorylation. The Src/FAK pathway can activate expression of certain matrix metalloproteinases, pointing to the importance of this pathway in cell motility^30, 44^. Thus, it is tempting to hypothesize that ZIP11 may also promote migration and invasion through activation of the Src/FAK pathway. Considering that ZIP13 deletion results in similar phenotypes to our observations on ZIP11 KD, the activation of Src/FAK kinases represent a viable mechanism to be investigated in order to understand the contribution of ZIP11 to cervical cancer cell metastasis.

However, our previous RNAseq analyses also showed that the expression of components of the Notch signaling pathway is also altered. The Notch pathway has been shown to play both an oncogenic and tumor suppressor role in a variety of cancer types^45^. For instance, the Notch1 receptor is required for cell fate determination, cell proliferation, differentiation, and apoptosis^46^. Activating mutations of *NOTCH1* render a constitutively active Notch signaling pathway, which ultimately leads to cancer^45, 47^. Over 80% of primary cervical cancer tumors showed altered expression of positive and negative regulators of the Notch pathway^47^. Active Notch1 has been shown to function in conjunction with papillomavirus oncogenes E6 and E7 in transformation of epithelial cells, which also resulted in resistance to anoikis, which is the programmed cell death that occurs during tumor metastasis^48, 49^.

Consistently, KD of *Notch1* in cervical cancer cells resulted in decreased activity of RhoC, a GTPase important in regulating cancer cell metastasis, and subsequently decreased cell migration and invasion^47, 50^. Given that *ZIP11* KD resulted in downregulation of *NOTCH1*^24^, it can be hypothesized that proper nuclear Zn transport is needed for expression and downstream activation of this signaling cascade, and therefore for the pro-oncogenic roles of activated Notch signaling in cervical cancer progression. Thus, the phenotypes dissected here in KD cells expressing the MBS mutations may be explained by alterations in nuclear Zn homeostasis which may result in reduced *NOTCH1* expression. Conversely, the functional phenotypes rendered by the predicted SNPs related to aggressive phenotypes evaluated here, are consistent with a role for these mutations in cervical cancer tumorigenesis and potentially rescuing the function of the Notch signaling pathway.

#### ZIP11 SNPs: the proximity to substrate binding region matters

When our lab first identified the ZIP11 SNPs in cervical cancer patients, A234P and P243S were predicted to have the strongest deleterious biological consequences^24^. These two SNPs were confirmed here to have the strongest ability to restore functional carcinogenic traits in HeLa cells. These two SNPs not only revived the proliferation defect in *ZIP11* KD cells, but also exhibited a robust expression and succeeded in bringing nuclear Zn levels back to normal, and consequently displayed similar migration and invasion capabilities compared to control cells. To better understand the phenotype of A234P and P243S, we turned to our *in-silico* alignment between hZIP11 and BbZIP4^18^ and the AlphaFold model. Both A234P and P243S align with specific residues, T201, and P210, located in TM5 of BbZIP4, respectively. Based in their position, it is plausible that these SNPs in fact produce changes in ZIP11 structure, which potentially increase substrate availability, as suggested by our previous report^24^. The key may be grounded in how these residues interact with surrounding helices and MBS in the core of the transmembrane domain and hydrophobic residues that block the metal transport pathway on the extracellular side^18^. We hypothesize that the A234P and P243S mutations could trigger electrostatic interactions among additional MBS, promoting metal release and rearrangement of the TMs, opening up the transporter on the cytosolic side. The substitution of alanine with proline in A234P would cause a structural bend or kink in the α-helix, potentially leading to a more accessible extracellular side for Zn. Further biochemical characterization of these transporters is undergoing and is beyond the biological scope of this manuscript.

While A234P and P243S proved to be powerful in their restorative abilities, the SNPs A26S and A89V showed a milder effect. The A89V maintained the phenotypic defects observed in KD cells, while the A26S successfully restored accumulated nuclear Zn levels and the migration defect but less effectively than A234P and P243S. However, the A89V mutation could restore the invasion defect, though not as efficiently. To better understand the subdued restorative effects of A26S and A89V we evaluated their position in the predicted ZIP11 structure. Unlike A234P and P243S, A26S and A89V were located on the outer helices of ZIP11, farther away from the substrate-binding region. BbZIP4, our reference, consisted of 8 TMs forming a helix bundle, with 4 inner helices surrounded by 4 outer ones^18^. The alignment and AlphaFold model hinted that A89V aligned with V137 on an outer TM3 of BbZIP4, while A26S aligned with A34, which was not part of any TMs constituting BbZIP4^18^. The intriguing part was that A89V showed accumulated nuclear Zn levels and slower migration rates compared to KD cells. We speculated that the replacement of alanine with valine, a hydrophobic residue, might have blocked the transporter’s extracellular side or hindered the helices’ conformational change required for metal release^18^. However, as BbZIP4 already had a valine at V137, the alanine-to-valine substitution might not significantly affect metal transport in either transporter. A detailed biochemical characterization is in progress to understand the details of the mechanism of transport *in vitro*.

Conservation of function of MBS between metal transporters and across species is well established. Consistently, in the alignment between hZIP11 and BbZIP4, three of the ZIP11 MBS aligned precisely with MBS residues in the proposed BbZIP4 ion transport pathway^18^. H204 in ZIP11 corresponded to H177 on TM4 of BbZIP4, a crucial player proposed to release metals from M1 of the binuclear metal center in BbZIP4. Following the path of the metal substrate, the E208 in hZIP11 aligned with E181 on TM4 of BbZIP4, while E244 in ZIP11 corresponded to E211 on TM5^18^. According to the published structure, the bridging residues, E181 and E211, are pivotal in connecting the two metal ions at M1 and M2 of the binuclear metal center, facilitating the transport and be released into the cytosol. Due to the conservation of these critical metal-binding residues between BbZIP4 and hZIP11, we propose that the MBS mutations may disrupt the formation of the binuclear metal center in ZIP11, leading to impaired metal transport and release. In addition, based on our western blot and confocal microscopy analyses, it seems like the alanine substitutions in H204 and E208 negatively impact protein stability, as their expression is significantly reduced than wild type and other mutant ZIP11 proteins expressed here. The position, decreased protein expression and biochemical relevance of these residues, provided a plausible explanation for the observed growth, migration, and invasion defects, and increased nuclear Zn levels, in *ZIP11* KD cells expressing these mutations. Importantly, KD cells expressing E244A displayed a partially restored proliferation defect, unlike the other MBS mutations. The location of E211 in BbZIP4, which corresponds to E244A in ZIP11 on the TM5, appeared to be the key to this behavior, and is potentially related to its proximity to the interphase between the membrane and cytosol. H204A and E208A, were located on TM4 in the middle of the transmembrane domain, a position that makes them essential for metal transport. Studies to characterize the biochemical properties of these proteins are also undergoing.

Overall, our study provides evidence linking specific ZIP11 SNPs and the relevance of metal transport to the establishment and maintenance of fundamental carcinogenic phenotypes in HeLa cells. Further research is needed to unravel the precise functions and regulatory mechanisms of Zn transporters in different cellular contexts and their implications in cancer biology and other diseases.

## MATERIALS AND METHODS

### Homology modeling and sequence analyses

Homology modeling and sequence analyses were performed using MUSCLE^51^ and ESPript 3.0^52^ software. The human ZIP11 isoform 1 sequence (NP_001153242.1) and BbZIP4^18^ (PDB: 6PG1) were used as references. The *predicted structure* of human ZIP11 (PDB AF-Q8N1S5-F1) from the AlphaFold Protein Structure Database^53^.

### Plasmids and site directed mutagenesis

The MISSION® pLKO.1-puro Non-Target shRNA (scr) and the shRNA against the UTR of human ZIP11 gene (5’-TCCTGATTGACTCTGATTATA-3’, Cat. TRCN0000434903) were from Sigma-Aldrich. The mammalian gene expression vector pLV[Exp]-EGFP/Neo-Ef1A (pLV) encoding human ZIP11 and empty vector were obtained from Vector Builder and previously reported^24^. pLV-ZIP11 was used as template DNA into which the individual SNPs and MBS mutations were introduced. The AccuPrime™ Pfx Supermix (Invitrogen, Cat. 12344-040) and primers listed in **Table 1** were used following the manufacturer’s protocol. Mutations were verified by Sanger sequencing.

**Table 1:**
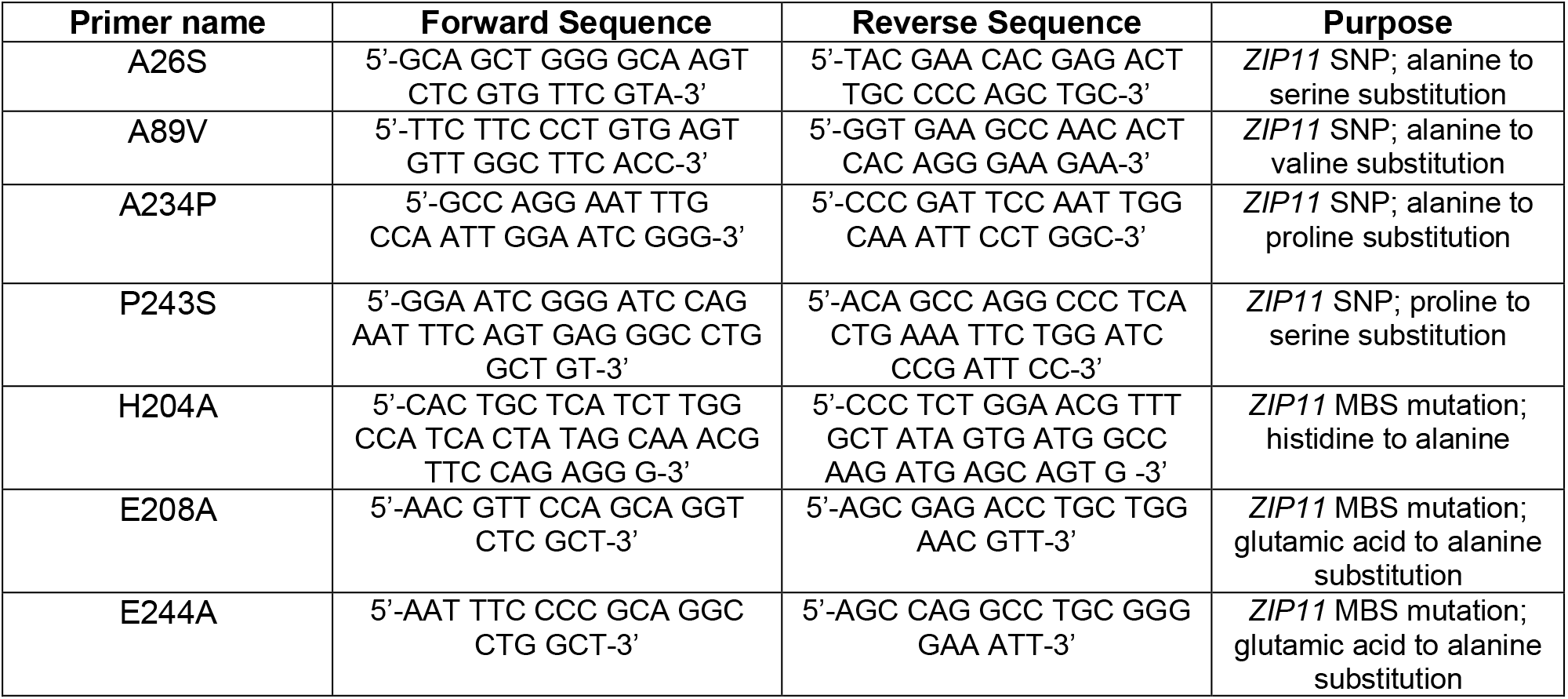
Primers used in this study.

### Cell Culture

The cervical cancer cell line HeLa and HEK293T fibroblasts were obtained from American Type Culture Collection (ATCC, Manassas, VA, United States). Stable KD HeLa cells for *ZIP11* were previously obtained and reported by our laboratory^24^. In the previous publication this *ZIP11* shRNA was named shRNA-2, and we continue to use this nomenclature for consistency. Cells were cultured in DMEM media (Sigma-Aldrich, St Louis, MO, United States) supplemented with 10% fetal bovine serum (FBS) and 1% antibiotics (penicillin G/ streptomycin, Gibco, Waltham, MA, United States) in a humidified atmosphere containing 5% CO_2_ at 37°C. Cells transduced with plasmids containing puromycin and gentamicin resistance cassettes were selected in media containing 2 µg/ml puromycin (Invitrogen, Waltham, MA, United States) or 4 mg/ml of geneticin, respectively. Then, cells were maintained in 1 µg/ml puromycin (Invitrogen) or 2 mg/ml of geneticin, respectively.

### Lentivirus Production and Transduction

Lentiviruses were produced in HEK293T cells using 15 µg of *ZIP11* vector encoding the SNPs or MBS mutations, 15 µg of the pLV1, 6 µg of pLV2, and 3 µg of pSVGV packing vectors using Lipofectamine 2000 (Thermo Fisher Scientific) following the manufacturer’s instructions. Lentiviral particles were collected from the media after 24 and 48 h post-transfection. The viral supernatant was filtered through a 0.22 µm syringe filter (Millipore Sigma, Burlington, MA, United States) and then used to transduce HeLa cells stably KD for *ZIP11* shRNA.

### Antibodies

The rabbit anti-ZIP11 (PA5-20679) was from Thermo Fisher Scientific (PA5-20679). The rabbit anti-GAPDH (A19056) was from Abclonal Technologies (Woburn, MA, United States). The mouse anti-MTF1 ((H-6): sc-365090) was from Santa Cruz Biotechnologies (Dallas, TX, United States). The secondary HRP-conjugated anti-mouse and anti-rabbit used for western blot were from Invitrogen (31430 and 31460, respectively). The fluorescent goat anti-rabbit secondary conjugated with Alexa 633 and a goat anti-mouse secondary antibody conjugated with Alexa 594 were acquired from Thermo Fisher Scientific (A-21070 and A-11032, respectively).

### Western Blot Analyses

Protein samples from HeLa cells including wildtype, scr shRNA control, *ZIP11*-KD, and cells transduced with the empty or ZIP11 SNPs and MBS mutants-containing pLV vectors were solubilized in RIPA buffer (10 mM piperazine-N,N-bis(2-ethanesulfonic acid), pH 7.4, 150 mM NaCl, 2 mM ethylenediamine-tetraacetic acid (EDTA), 1% Triton X-100, 0.5% sodium deoxycholate and 10% glycerol) supplemented with Complete protease inhibitor cocktail (Thermo Fisher Scientific). Protein content was quantified by Bradford assay^54^ and 20 μg of protein of each cell line were separated by SDS-PAGE and electrotransferred to PVDF membranes (Millipore Sigma). The primary antibodies anti-ZIP11 and anti-GAPDH as a loading control, were incubated overnight at 4 °C followed by a 2 h incubation of the species-specific secondary antibodies coupled to horseradish peroxidase. Membranes were developed using the high sensitivity Tanon reagent (Abclonal Technologies).

### Confocal Microscopy

HeLa cells were cultured to 100% confluency and were fixed in 10% formalin-PBS overnight at 4 °C. The cells were washed with PBS and permeabilized with 0.2% Triton X-100 in PBS for 15 min and incubated in blocking solution (PBS, 0.2% Triton X-100, 3% FBS) for 1 h at room temperature. Samples were incubated with the primary antibodies rabbit anti-ZIP11 or mouse anti-MTF1, in blocking buffer overnight at 4 °C. The following day the samples were incubated with fluorescent goat anti-mouse Alexa-594 and anti-rabbit Alexa-633 secondary antibodies in blocking solution for 2 h at room temperature. DAPI was used to stain the nuclei and samples were mounted with VectaShield mounting solution (H-1200; Vector Laboratories, Inc, Newark, CA, United States). Confocal microscopy was performed on a Leica SP8 images were analyzed with the Leica Application Suite X (Leica Microsystems Inc., Buffalo Grove, IL, United States). The pLV vector also encodes a fluorescent green fluorescent protein (GFP) tag, the GFP level of each strain was checked periodically to verify constitutive expression of the transduced DNA.

### Cell Proliferation Assays

HeLa cells were seeded at a density of 1 x 10^4^ cells/cm^2^. Samples were collected 24, 48, 72, and 96 h after plating. Cells were washed with 200 µl of PBS, trypsinized, then counted using a Cellometer Spectrum (Nexelcom Biosciences, Lawrence, MA, United States). To assess Zn resistance of HeLa cells overexpressing the SNPs and MBS constructs, each stable cell line was cultured with increasing concentrations of Zn as described before^24^. Then, cells were counted at 72 h, as it was a timepoint where metal sensitivity was evident in previous studies^24^.

### Metal Content Analysis

HeLa cells were seeded at a density of 1 x 10^4^ cells/cm^2^ and grown until they reached 100% confluency. The Rapid, Efficient, and Practical (REAP) nuclear and cytoplasmic separation method was used to isolate subcellular fractions, and samples were analyzed by atomic absorbance spectroscopy (AAS), as previously described^24, 55–57^. Briefly, the cells were washed three times with ice-cold PBS free of Ca^2+^ and Mg^2+^ (Gibco). The cells were resuspended and lysed in 500 µl of ice-cold PBS containing 0.1% NP-40 (Sigma-Aldrich). The whole cell fraction (100 µl) was collected, and the remaining samples were centrifuged for 10 s at 10,000 rpm. The supernatant was collected as the cytosolic fraction. The nuclear pellet was washed in 200 µl of the ice-cold PBS/ 0.1% NP-40 buffer and centrifuged for an additional 10 s. The nuclear pellet was resuspended in 200 µl of ultrapure water. Samples were sonicated for 5 min in cycles of 30 s on, 30 s off using a Bioruptor Plus Sonication System (Diagenode, Denville, NJ, United States). Protein was quantified by Bradford^54^ and samples were mineralized in trace metal grade nitric acid HNO_3_ and boiled at 100 °C.

Zinc levels were measured by AAS, using a PinAAcle 900Z AA Spectrometer (PerkinElmer) with Zinc (Zn) Lumina Hollow Cathode Lamp (PerkinElmer) at 213.86 nm wavelength. Zn standard solutions were prepared from 1000 mg/L (Sigma-Aldrich), and the matrix modifier in working solution was Mg(NO_3_) 1%v/v (PerkinElmer) in HNO_3_ trace metal grade (Thermo Fisher Scientific). The linearity considered for the measurements 2.5 - 20 ppb, with an R2 >0.99 carried out at least in triplicates. When required, samples were diluted with H_2_O (18 ΩM). Zn content on each sample was normalized to the initial protein content in each sample.

### Wound Healing Assay

HeLa cells were cultured in 6 well plates until they reached 100% confluence. Cells were then starved in DMEM without FBS for 24 h and treated with Cytosine β-D-Arabinofuranoside (AraC, Thermo Fisher Scientific) for 2 h to block cell proliferation. Then, the monolayer was scratched with a sterile 200 µl pipette tip and washed with PBS to remove floating cells. Closure of the wound was imaged every 24 h until wound closure. Regions of the wound were marked on the plate so that pictures could be taken in a consistent location. FIJI software, version 1.44p, was used to quantify the area migrated by the cells into the wound at each time point^58^.

### Matrigel Invasion Assay

The Transwell chamber method was used to measure the invasive capabilities of each strain of HeLa cells overexpressing wild type ZIP11 and SNPs and MBS mutations. HeLa cells were treated with AraC for 2 h to block proliferation and plated at a density of 1.25 x 10^5^ in 2 ml of DMEM without serum in the upper chamber of the BioCoat® Matrigel® Invasion Chambers (Corning), and DMEM supplemented with 10% FBS served as the chemoattractant in the lower chamber as previously described^24, 30^. Cells were incubated for 24 h at 37 °C in a 5% CO_2_ atmosphere and cells that had invaded through the Matrigel and membrane were fixed in methanol for 5 min, stained with 0.1% crystal violet diluted in PBS. Ten images of three independent biological replicates were taken with an Echo Rebel microscope (San Diego, CA, United States) and analyzed using the FIJI software (version 1.44p)^58^. The invasion index was calculated as the ratio between number of cells of wild type and *ZIP11* KD cells controls transduced or not with scr shRNA, pLV empty vector, KDs reconstituted with EV or wild type ZIP11 and the SNPs and MBS mutants and the number of WT control cells.

## ACKNOWLEDGEMENTS

We thank Dr. José Argüello’s laboratory at Worcester Polytechnic Institute for facilitating access to their Atomic Absorption Spectrometer. The authors are also thankful to Mr. Jaime Carrazco-Carrillo and Mr. Michael Quinteros for their technical assistance.

## AUTHOR CONTRIBUTIONS

TP-B conceived and designed the research; EK, OV-T, KD-R, FH, and TP-B performed experiments and compiled data; EK, KD-R, LC and TP-B analyzed data; EK and TP-B prepared figures and tables; EK, FH and TP-B drafted the manuscript; all authors edited and revised the manuscript; all authors approved the final version of the manuscript.

## CONFLICT OF INTEREST

The authors declare no conflicts of interest.

### Data availability

The data original for this study are available in the manuscript. Additional raw data and replicate data are available and will be shared on reasonable request to the corresponding author.

RNAseq analyses referred to in this work are published^24^ and available at GEO with the accession number GSE198411.

